# Combined Central and Peripheral Nerve Stimulation Improves Functional Recovery of Mixed Peripheral Nerve Injury in a Rat Forelimb Model

**DOI:** 10.1101/2025.01.26.634922

**Authors:** Sahand C. Eftekari, Yong-Chul Yoon, Rex Chin-Hao Chen, Peter J. Nicksic, D’Andrea T. Donnelly, Anna Jesch, Ellen C. Shaffrey, Weifeng Zeng, Kip A. Ludwig, Samuel O. Poore, Benjamin J. Vakoc, Aaron J. Suminski, Aaron M. Dingle

**Affiliations:** University of Wisconsin School of Medicine and Public Health, Division of Plastic Surgery, Madison, Wisconsin, USA; Massachusetts General Hospital, Wellman Center for Photomedicine, Boston, Massachusetts, USA; University of Wisconsin School of Medicine and Public Health, Department of Neurosurgery, Madison, Wisconsin, USA

**Author notes:** **Correspondence to:** Aaron M. Dingle PhD, University of Wisconsin School of Medicine and Public Health, Division of Plastic Surgery, 1111 Highland Avenue, 5107 Wisconsin Institutes for Medical Research. Equal contributions (Aaron J. Suminski, Aaron M. Dingle).

**Keywords:** electrical stimulation, nerve regeneration, trigeminal nerve, nerve injury

## Abstract

**Introduction:** Peripheral nerve reinnervation following nerve injury is often a slow and incomplete process, resulting in significant morbidity and permanent loss of function of the injured extremity in many patients. Prior studies have shown the efficacy of electrical stimulation to synchronize the axonal regeneration of both motor and sensory neurons in peripheral nerve injury models. Moreover, separate investigations have also shown the use of cranial nerve stimulation, principally the vagus nerve, to improve functional outcomes. However, no study has investigated the synergistic effects of both intraoperative electrical stimulation and cranial nerve stimulation for functional improvement within a peripheral nerve injury model. This investigation quantifies the efficacy of combined intraoperative electrical stimulation and trigeminal nerve stimulation on motor and sensory functional recovery in a rat peripheral nerve injury model.

**Methods:** Twelve adult male Lewis rats were trained in a reach and pull task for a food reward using their right forelimb with baseline force thresholds and percent success of the pull task recorded. Baseline sensory data was retrieved using an automated von Frey monofilament test. All rats underwent surgical transection and 2mm gap repair of their median and ulnar nerve of their right forelimb followed by 1 hour of intraoperative electrical stimulation. Trigeminal nerve stimulation throughout the rehabilitation period was completed via supraorbital nerve headcap electrodes. Motor and sensory data were compared to historic cohorts comprised of sham surgery (no nerve injury), brief intraoperative electrical stimulation, trigeminal nerve stimulation, and a sham peripheral and trigeminal nerve stimulation group. Polarization sensitive optical computed tomography (PS-OCT) was used to assess nerve regeneration in fixed tissue samples.

**Results:** The combined cohort of rodents were able to recover to their pre-injury motor function by the third week of rehabilitation, faster than either of the singular electrical stimulation cohorts assessed previously. Moreover, functional sensory data of the combined stim cohort demonstrated no change when compared to their pre-injury baseline’.

**Conclusions:** Peripheral nerve electrical stimulation and trigeminal nerve stimulation are two separately acting mechanisms of therapy that employ electric waveforms to improve the functional recovery of injured peripheral nerves. The former acts within the periphery to synchronize axonal growth and regeneration of the injured neurons, while the latter acts centrally to augment neuroplasticity. When used simultaneously in a rodent peripheral nerve injury model, these modalities have shown to build upon each other to deliver a faster functional motor recovery, while sensory recovery outcomes remain to be demonstrated.

## Introduction

Limb trauma is a common affliction in the United States, with an estimated 300,000 patients per year presenting to the emergency department with this chief concern.^1^ Of these limb trauma patients, 1.64% have an associated peripheral nerve injury.^1^ This leads to more than 50,000 annual peripheral nerve repair procedures in the United States alone.^2^ Unfortunately, peripheral nerve reinnervation is often a slow and incomplete process, resulting in significant morbidity and permanent loss of function of the injured extremities in approximately 50% of patients even with surgical intervention.^3^ These injuries have the potential to create devastating deficits in a patient’s life and work. Therefore, innovative methods to augment peripheral nerve regeneration and optimize functional outcomes are needed.

Electrical stimulation at the site of the peripheral nerve injury has been demonstrated to improve the functional outcomes of injured nerves.^2,4,5^ It is thought that this stimulus induces a synchronized depolarization across all the afflicted neurons and their supporting Schwann cells to trigger the regeneration of the injured nerve at one time point.^6^ This peripheral stimulation (PS) therapy has shown efficacy in clinical trials as well, with the BEST SPIN randomized control trial showing marked improvement of shoulder function with intraoperative electrical stimulation of the spinal accessory nerve following neck dissection and injury to the nerve.^6^

More recently, and through a separate and largely speculated mechanism, studies have shown the use of cranial nerve electrical stimulation to improve the functional outcomes of peripheral nerve injury. Stimulating the afferent neurons of a cranial nerve is thought to induce a larger neuronal circuit that begins at the brainstem and projects up to the cerebral cortex to promote neuroplasticity of a desired cortical region.^4,5^ During neuronal regeneration after peripheral nerve injury, many axons will inevitably grow into incorrect nerve tubes and innervate inappropriate motor and sensory targets. The brain must then relearn these new connections, requiring rewiring of the synapses of the associated cortical region.^7^ The use of cranial nerve stimulation in the setting of peripheral nerve injury has been studied using the repetitive stimulation of the vagus nerve coupled to a learned task in a rodent model.^7,8^ These research endeavors illustrate the increased synaptic reorganization and improved sensorimotor recovery modulated by stimulating the vagus nerve without any observable peripheral changes at the site of injury.^8^ Unfortunately, the anatomic course of the vagus nerve in humans requires the surgical implantation of a cuff electrode deep within the neck, and therapeutic electrical stimulation is stifled by adverse effects such as vocal cord spasm.^9,10^

An attractive alternative to vagal nerve stimulation is trigeminal nerve stimulation (TNS). This nerve courses much more superficially within humans and has the capability to be stimulated through non-invasive methods such as transcutaneous electrical stimulation (TENS) units.^11,12^ The afferent neurons of the trigeminal nerve also allow access to deep brain nuclei that provide gateways for triggering larger projections of electrical activity in the cerebral cortex and can thus provide a similar window for central nervous system stimulation.^10,13^

A recent investigation within our laboratory conducted by Nicksic et al. has quantitatively compared the functional recovery of injured peripheral nerves using either PS or TNS compared to no electrical stimulation in a rodent model.^14^ This investigation found that rodents within the PS group were able to return to baseline motor function after 4 weeks of rehabilitation, while rodents within the TNS group were able to return to baseline motor function after 6 weeks of rehabilitation, both faster than the no electrical stimulation cohort.^14^ Moreover, functional sensory recovery of these rodents was observed to return to baseline after 3 and 1 weeks of rehabilitation in the PS and TNS groups, respectively.^14^

Although the Nicksic et al. investigation was able to produce quantifiable data on the functional improvement of motor and sensory recovery of peripheral nerves using either PS or TNS, no cohort of rat received *both* PS and TNS as a combined therapy.^14^ Given that these mechanisms are thought to augment peripheral nerve regeneration through separate and mutually exclusive mechanisms, we hypothesized that these mechanisms can be summated to provide a combined further improvement in peripheral nerve regeneration. Our investigation aims to utilize a 5^th^ cohort of rodents receiving *both* PS and TNS under the same protocol and instruments. In doing so, we aim to quantify the effects of combined PS and TNS on the functional motor and sensory recovery of peripheral nerves in a rodent model and to compare these outcomes to the previous four cohorts (Figure 1). Moreover, we aim to histologically analyze the regenerated nerve cross sections for fascicular organization and scarring. Our laboratory decided to pursue a 5^th^ cohort rather than re-instituting an entirely new investigation due to concerns of rodent attrition during peripheral nerve injury surgery and the ethical considerations to minimize the number of animals within our research.

**Figure 1:**
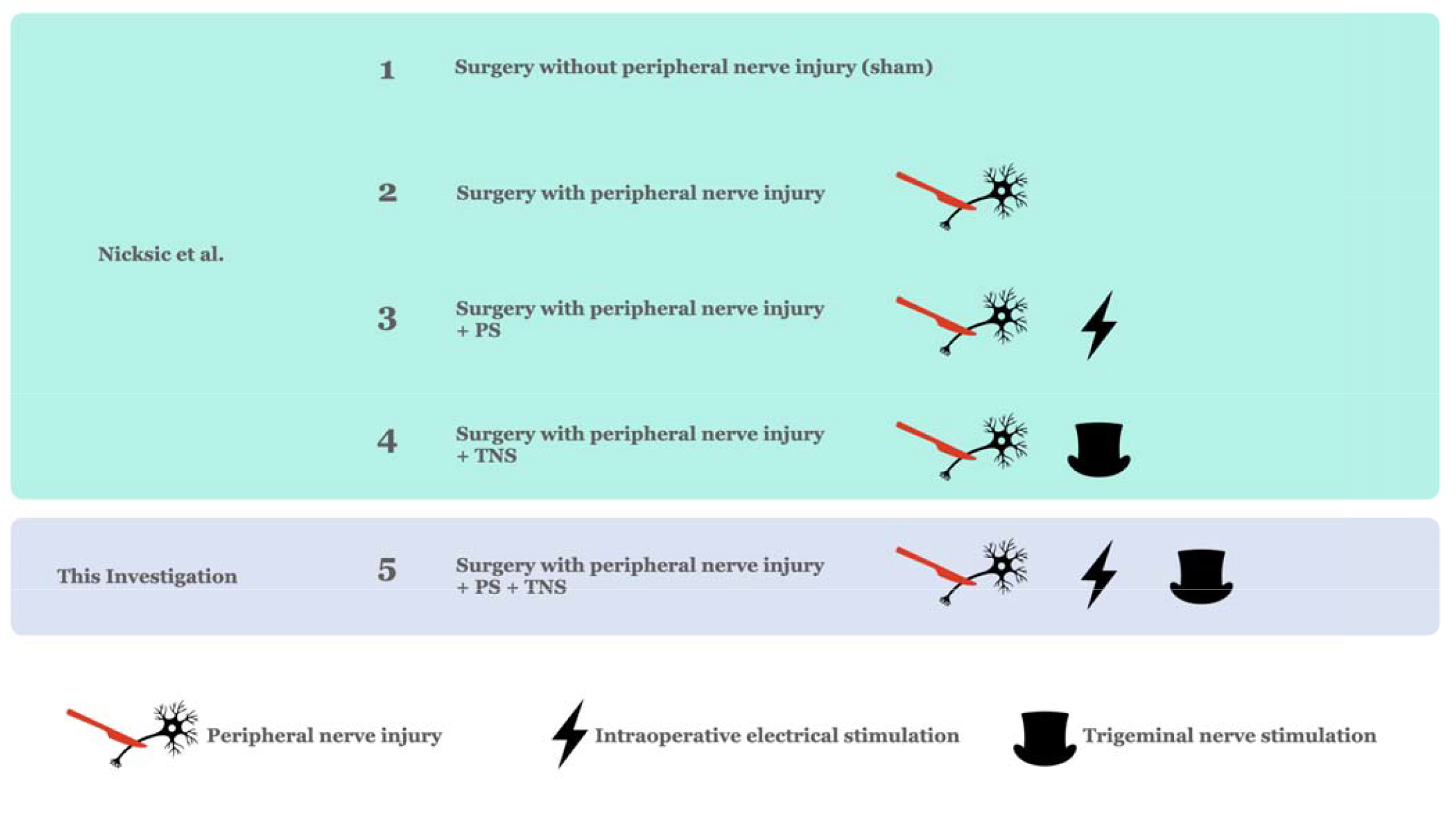
Comparison of rodent cohorts in the work of Nicksic et al. and the addition of a fifth cohort to assess the combined effect of peripheral stimulation (PS) and trigeminal nerve stimulation (TNS).^14^

**Figure 2:**
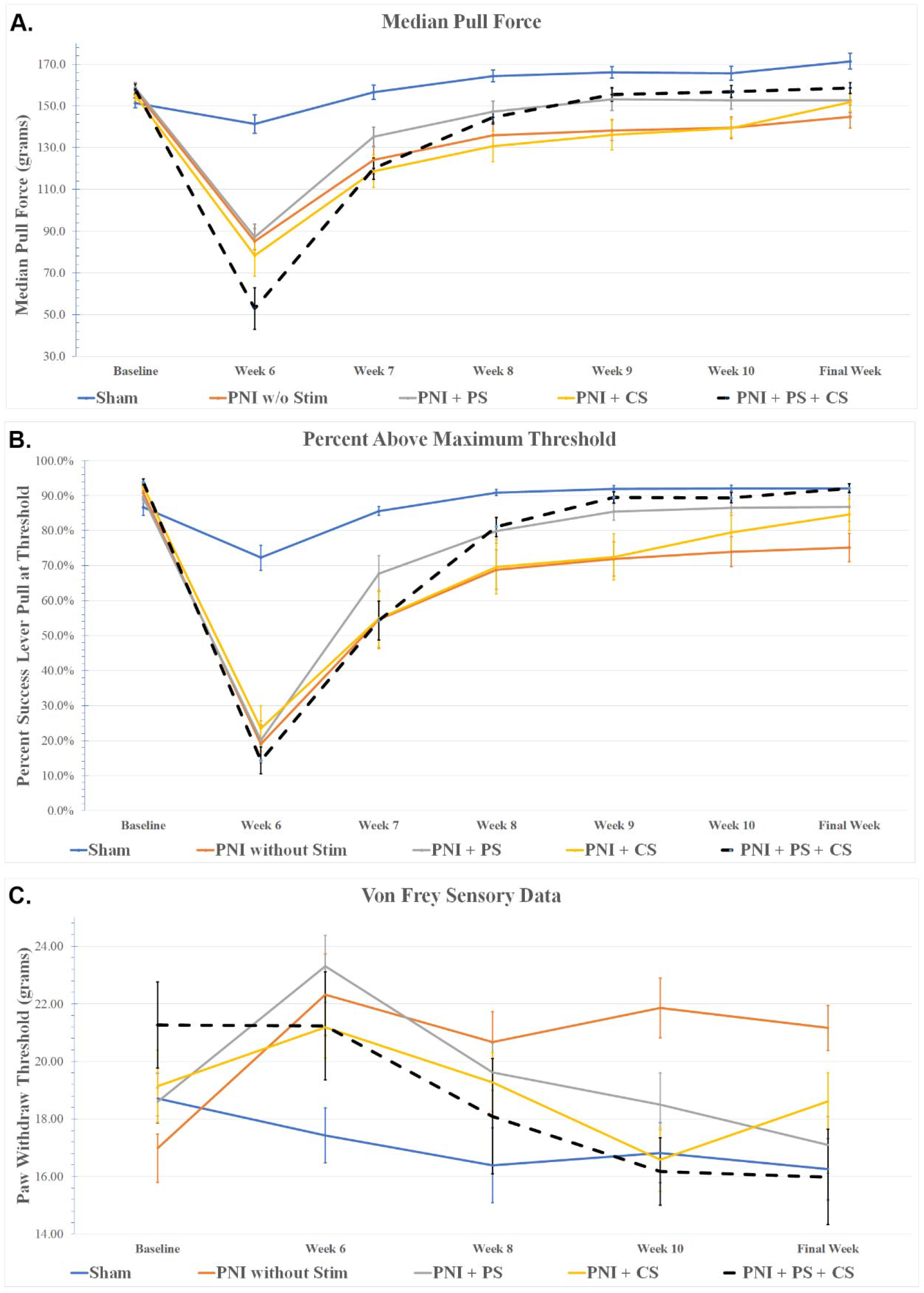
**A-C:** A. Median pull force, B. Percent success. C. Sensory paw withdrawal thresholds of PS+TNS cohort compared to historic data for sham surgery, Peripheral nerve injury (PNI) without stim, PNI with peripheral stimulation (PS), PNI with trigeminal nerve stimulation (TNS), and PNI with both PS and TNS.^14^

## Methods

Twelve adult male Lewis rats weighing approximately 250g were included in this study, mirroring the cohorts from previous work.^14^ Lewis rats were selected due to their low rate of denervation-related autophagy and their demonstrated ability to be trained for a food-rewarded behavior.^14, 15^ All aspects of the protocol were approved by the University of Wisconsin Institutional Animal Care and Use Committee, designed in accordance with ARRIVE guidelines, and all methods were performed in accordance with the relevant guidelines and regulations. All animals were housed in individual cages in the vivarium with a 12-hour light cycle environment. All regulations, protocols, equipment, and instruments used were the same as the Nicksic et al. investigation, with the exception of a new lead investigator (SCE). For further details of the subsequent methodology, refer to the works of Nicksic and colleagues.^14^

### Baseline Assessments

All rodents were trained to proficiency in an isometric reach and pull task with their right forelimb using the MotoTrak system (Vulintus, Inc., Dallas, TX) over a 6-week period for a food reward (45 mg dustless chocolate precision pellet, BioServe, Frenchtown, NJ, USA).^8^ Baseline sensory data was collected using a von Frey monofilament (Ugo Basile, Barese, Italy) sensory assessment device.^16^

### Initial Injury Surgery

Following baseline training and assessment, all rodents underwent surgery on their right forelimb for sharp transection of their median and ulnar nerve followed by a 2mm microsurgical coaptation using a silastic cuff. A custom fabricated bipolar nerve cuff was fastened proximal to the coaptation site for one hour of continuous electrical stimulation of the median and ulnar nerve *en bloc*. Electrical stimulation parameters were identical to those previously described and based on prior literature (biphasic, cathode leading, charged-balanced pulses; 20 Hz low frequency; 100 μs pulse width).^14^ Following surgery, all subjects were allotted 5 weeks for recovery in the vivarium with unrestricted food intake. The contralateral uninjured limb served as a control.

### Trigeminal Nerve Cuff Implantation

After the initial recovery period, all subjects underwent a second surgery for implantation of a trigeminal nerve electrode cuff to the left supraorbital nerve to enable trigeminal nerve electrical stimulation contralateral to the injured limb. This electrode was custom fabricated to create a headcap screwed into the subject’s calvarium. Four screws were placed between the coronal and lambdoid sutures of the rodent cranium. A custom-fabricated bipolar nerve cuff (0.6 mm internal diameter) was placed around the left supraorbital nerve. Intraoperatively, the rodents each had electrodes validated by eliciting a blink reflex as previously described.^11^ Moreover, impedance tested was performed during the electrode cuff placement and at weekly intervals during functional rehabilitation to ensure proper contact and dosage of stimulation of the supraorbital nerve. Following trigeminal nerve cuff implantation, all subjects were allotted 1 week for recovery in the vivarium with unrestricted food intake.

### Rehabilitation

Following recovery of the trigeminal nerve implantation surgery, all subjects returned to the MotoTrak system for rehabilitation of their reach and pull task for 6 weeks. The same protocol was employed throughout the rehabilitation as the training period. Metrics of median peak force of lever pulling and percent success at reaching minimum threshold were recorded. All animals were tethered by their headcap to a commutator throughout the rehabilitation period and received electrical stimulation for each successful lever interaction. The current of the trigeminal nerve electrode was titrated identical to previous work.^14^ Subjects were tethered by their headcap to a he commutator during the first 5 weeks of rehabilitation and were stimulated automatically each time they had a successful lever interaction (constant current, biphasic, cathode leading, charge-balanced pulses, 30 Hz; 200 μs pulse width; 500 ms pulse duration).^14^ Biweekly sensory data was obtained throughout the rehabilitation phase using von Frey monofilament sensory assessment device.^16^ In the final week of rehabilitation training, all subjects were allowed to complete MotoTrk sessions untethered to the headcap electrode wire.

### Euthanasia and Histology

Upon completion of the rehabilitation period, all subjects were anesthetized with isoflurane and euthanized via intracardiac injection of pentobarbital. At this time the subjects underwent resection of their median and ulnar nerves on both the right forelimb as well as healthy contralateral forelimb for histological preparation using Gomori’s trichrome stain.

### PS-OCT Nerve Imaging

PS-OCT imaging was performed on fixed nerve samples from the new and historic cohorts using a custom-designed system based on a design described previously^19-22^. The light source provided 110 nm bandwidth (HSL-200-50LC, Santec Corp., Japan) centered at 1310nm and operated with a 50 kHz A-scan rate. Axial resolution was measured to be 7 μm (in air). The PS-OCT system used a polarization-modulation scheme that switched sample-arm polarization launch states on even and odd A-lines^18^ using an electro-optic modulator (Boston Applied Technologies, United States). The OCT receiver supported polarization-diverse detection, and a linear polarizer was placed at the end of the reference arm to reduce noise induced by polarization-mode dispersion (PMD)^23-24^. The system provided X mW power to the sample. Imaging was performed using a 2-axis galvanometer microscope using a Thorlabs LSM02 objective. PS-OCT image processing followed methods previously described previously^19^. Briefly, the system calculated the depth-resolved Stokes data. Using these data, local birefringence and the axis of orientation was calculated, using spectral-binning to mitigate noise induced by PMD^25^. The axis of orientation data was mapped to a 2D plane using a principal analysis reduction described previously^19^, and the angular range was mapped to a cyclical hue colormap as shown in figures. The birefringence was scaled to lie between 0 and 1, and mapped to the value parameter of the HSV colormap. The absolute orientation of the axis of orientation data was not controlled, and as such the relative changes in color within a dataset are meaningful but the color differences from dataset to dataset are likely attributable to changes in the sample arm optical fiber birefringence. PS-OCT datasets were generated and visualized using imageJ. Cross-sectional images as well as volumetric projections were generated.

### Statistical Considerations

Twelve subjects were justified by power analysis (statistical power = 0.8) and previous experience^14^. Statistical methods of comparison were determined prior to the start of data collection. In order to compare motor and sensory data between groups, a repeated measures analysis of variance (ANOVA) was employed. The averaged median peak force, percent success of lever pull, and paw withdrawal thresholds for each week were compared among groups and across weeks. Post-hoc analysis was performed using the differences of least square means. A threshold significance of 0.05 for type I error was set for all statistical tests. The subjects were determined to reach functional recovery for each of the three metrics when the aggregated average value of the cohort for a given week was not statistically different than their pre-injury baseline. All values for functional motor and sensory outcomes are represented as mean differences from the pre-injury baseline ± standard error. A positive or negative mean difference denotes a value that was greater than or less than the pre-injury baseline value, respectively. Histological analysis quantitatively compared the ratio of fascicular volume to full volume of the nerve cross sections that were injured to the healthy left forelimb. Qualitative assessment of fascicular organization and scarring was also completed.

## Results

### Pre-Injury Baseline Functionality and Injury Severity Metrics

The PS+TNS cohort averaged a median pull force of 158.05±2.5g, a percent success of 93.9±0.9%, and a paw withdrawal threshold of 21.26±1.5g at baseline prior to injury surgery. There was no statistical difference in these metrics when compared to the previous four cohorts. Following initial injury surgery, the PS+TNS cohort demonstrated a 103.42±5.9g drop in median pull force and a 78.29±5.0% drop in percent success, indicating a significant decrease in motor functionality (P < 0.0001). All previous cohorts receiving peripheral nerve injury also demonstrated a statistically significant decrease in motor functionality (P< 0.0001). There was a 0.39±1.54g increase in paw withdrawal threshold which did not show a statistically significant decrease in sensory functionality (P > 0.05). This was inconsistent with the previous cohorts of that received peripheral nerve injury, which demonstrated a statistically significant decrease in sensory functionality following initial injury surgery.

At the beginning of the rehabilitation period, median pull force data of the PS+TNS cohort demonstrated a statistically significant lower motor function of the rodents compared to the sham surgery, no stim, and PS cohorts (P < 0.05). When compared to the TNS cohort, the PS+TNS cohort did not show any significant difference in median pull force performance (P > 0.05). The percent success of the PS+TNS cohort did not show any significant difference when compared to the no stim, PS, and TNS cohorts (P > 0.05) but did show a significant difference compared to the sham surgery cohort (P < 0.0001). There was no significant difference in sensory function among any of the cohorts at this time point (P > 0.05).

### Motor Recovery

The PS+TNS cohort was able to recover to its pre-injury median pull force and percent success performance by the third week of rehabilitation (week 8 post-injury) and demonstrate a mean difference of 1.85±5.86g pull force and 11.24±4.9% percent success mean difference at this time point (P > 0.05) (Figure 3).

**Figure 3:**
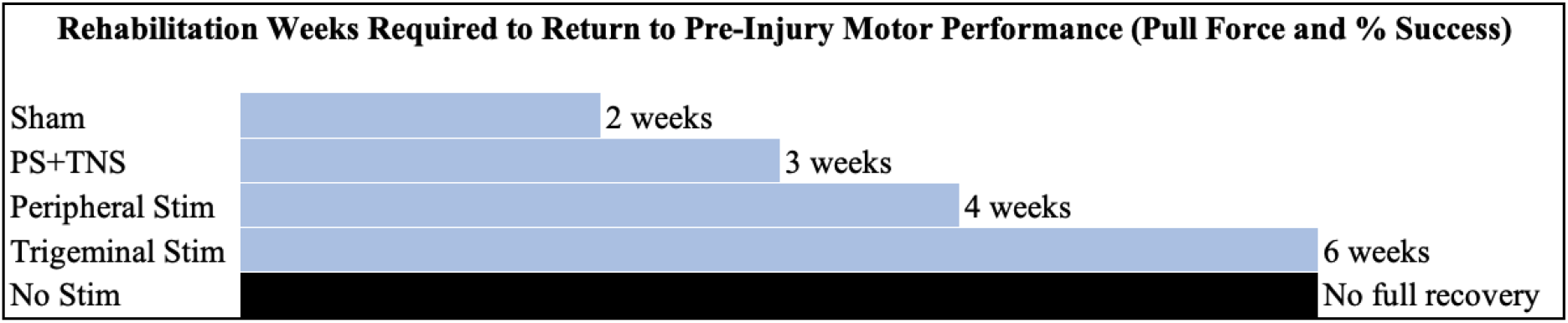
Rehabilitation time required to reach pre-injury functional motor performance.

The no stim cohort did not return to its pre-injury baseline for pull force (mean difference 12.36±5.48g, P = 0.0252) or percent success (mean difference 15.54±4.78%, P = 0.0013) by the final week of rehabilitation. The PS cohort recovered to its pre-injury median pull force performance (mean difference 5.38±5.25g, P > 0.05) and percent success (mean difference 3.99±4.58%, P > 0.05) at the fourth week of rehabilitation (week 9 post-injury). The TNS cohort returned to its pre-injury baseline of pull force (mean difference 3.873±5.48g, P > 0.05) and percent success (mean difference 18.51±4.78%, P > 0.05) at the final week of rehabilitation (week 11 post injury). Lastly, the sham surgery returned to its pre-injury baseline for median peak force (mean difference 22.19±6.06g, P > 0.05) and percent success (mean difference 1.311±5.29%, P > 0.05) at the second week of rehabilitation (week 7 post injury).

### Sensory Recovery

The PS+TNS cohort did not show a significant difference from pre-injury paw withdrawal to the beginning of rehabilitation (P > 0.05). Moreover, this was inconsistent with the previous four cohorts of rodents, which all demonstrated significant decrease in sensory function. All subsequent timepoints of paw withdrawal thresholds throughout the rehabilitation period of PS+TNS also did not show any statistical difference from baseline pre-injury assessment (P > 0.05) (**Figure 4**).

**Figure 4:**
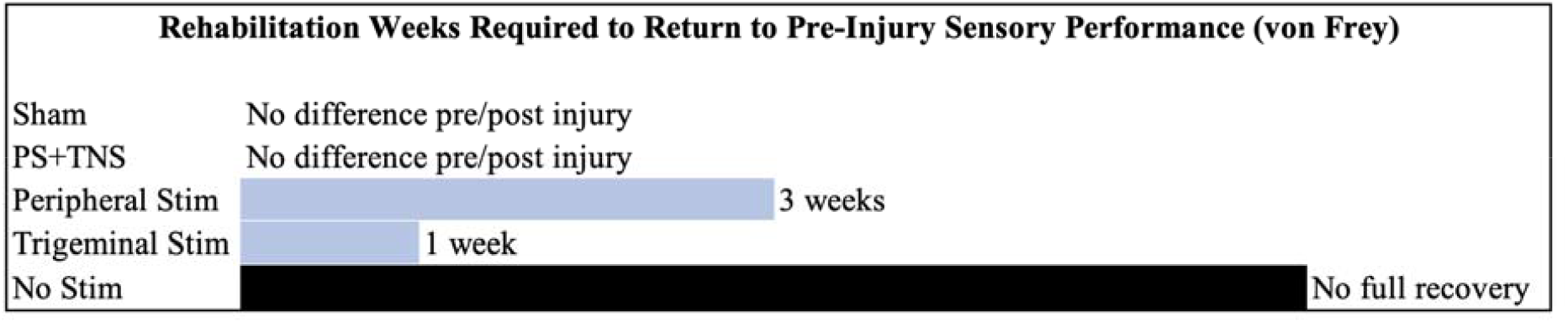
Rehabilitation time required to reach pre-injury functional sensory performance as measured with von Frey paw withdrawal thresholds.

The no stim cohort did not recover to the pre-injury baseline throughout the rehabilitation period for paw withdrawal (mean difference 4.18±1.59g, P = 0.0092). The PS cohort returned to its pre-injury baseline at the third week of rehabilitation (week 8 post injury) (mean difference 1.019±1.52g, P > 0.05) whereas the TNS group returned to baseline pre-injury sensory function at the first week of rehabilitation (week 6 post injury) (mean difference 1.08±1.59g, P > 0.05). The sham cohort demonstrated no difference in the pre-injury paw withdrawal thresholds compared to any timepoint throughout the rehabilitation period (P > 0.05).

### Nerve Histology

There was no statistically significant difference in the histomorphological examination of injured (right) and uninjured (left) nerves. The right median (RM) and right ulnar (RU) nerve cross sectional fascicular area to collagen ratio of the transected nerves (RM 0.578±0.096, RU 0.628±0.108) compared to the left median (LM) and left ulnar (LU) healthy nerve ratios (LM 0.601±0.065, LU 0.625±0.110) (P > 0.05) (**Figure 5**). Qualitative assessment of the injured nerves demonstrated substantially more epineural collagen deposition around the nerve cross section compared to the healthy counterparts (**Figure 6**).

**Figure 5:**
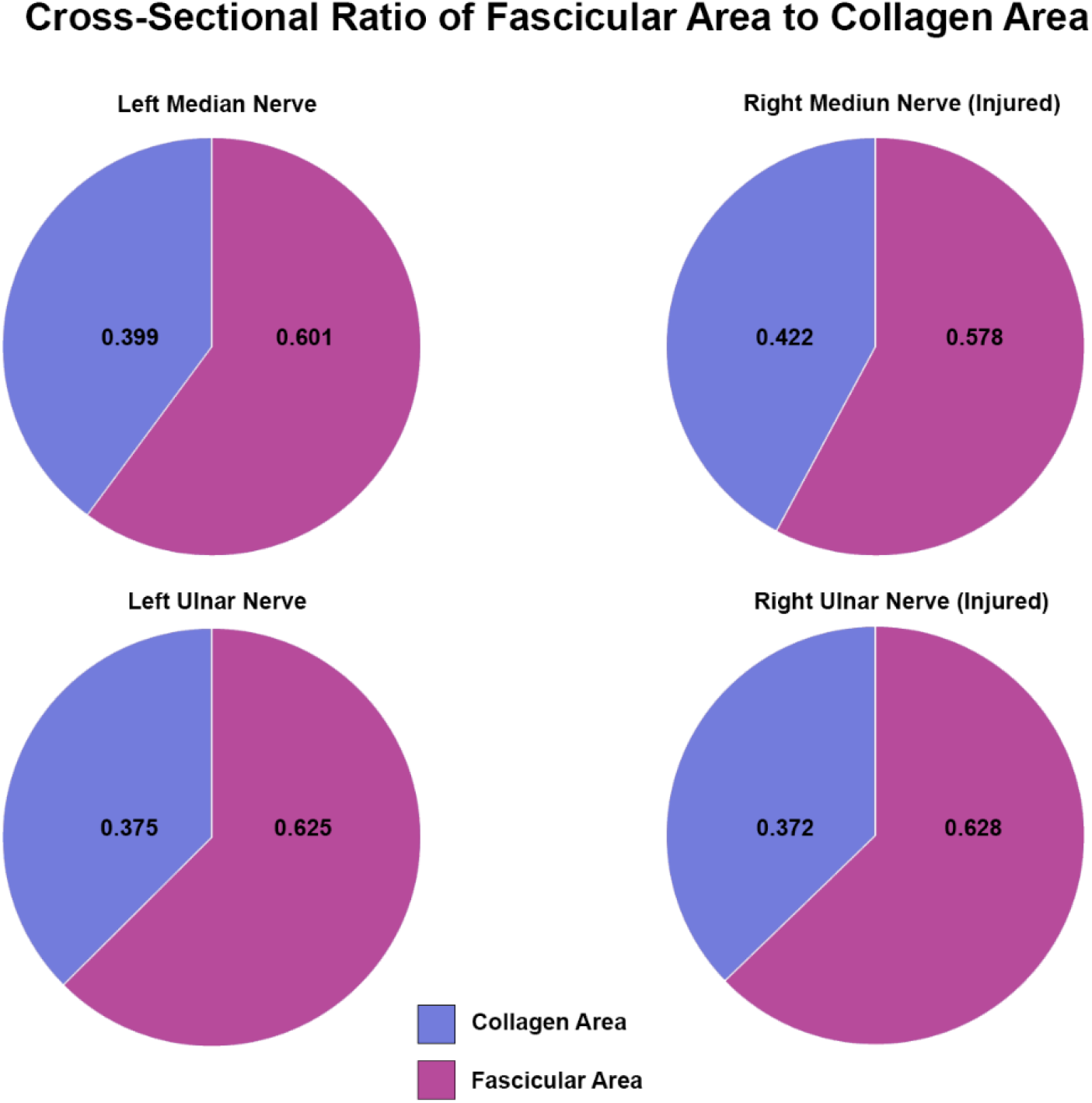
Ratio of the right transected and regenerated median and ulnar nerve fascicular area to collagen area compared to the healthy median and ulnar nerve ratios. No statistical difference exists among the groups.

**Figure 6:**
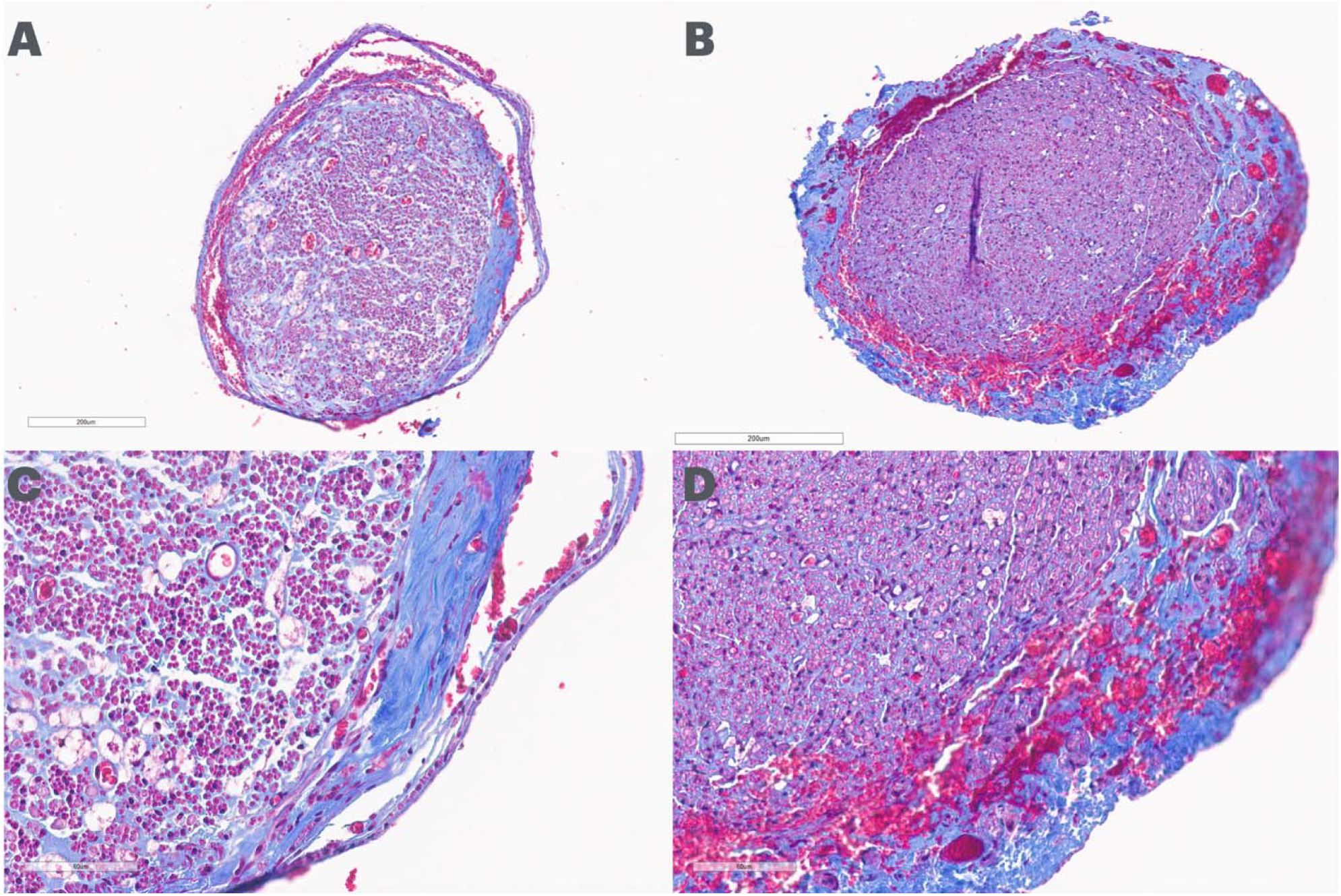
Representative nerve histomorphology of uninjured (A & C), and injured (B & D) median nerves. Qualitatively, injured nerves demonstrate a more dense fascicular area with amore disorganized and thicker ring of epineurial collagen. **Transverse nerve sections stained with Gomori’s trichrome; which stains myelinated axons purple and collagen blue, imaged at low (A & B: 11.5x magnification, scale bar = 200μm) and high (C & D: 40x magnification, scale bar = 60 μm)**.

**Figure 7:**
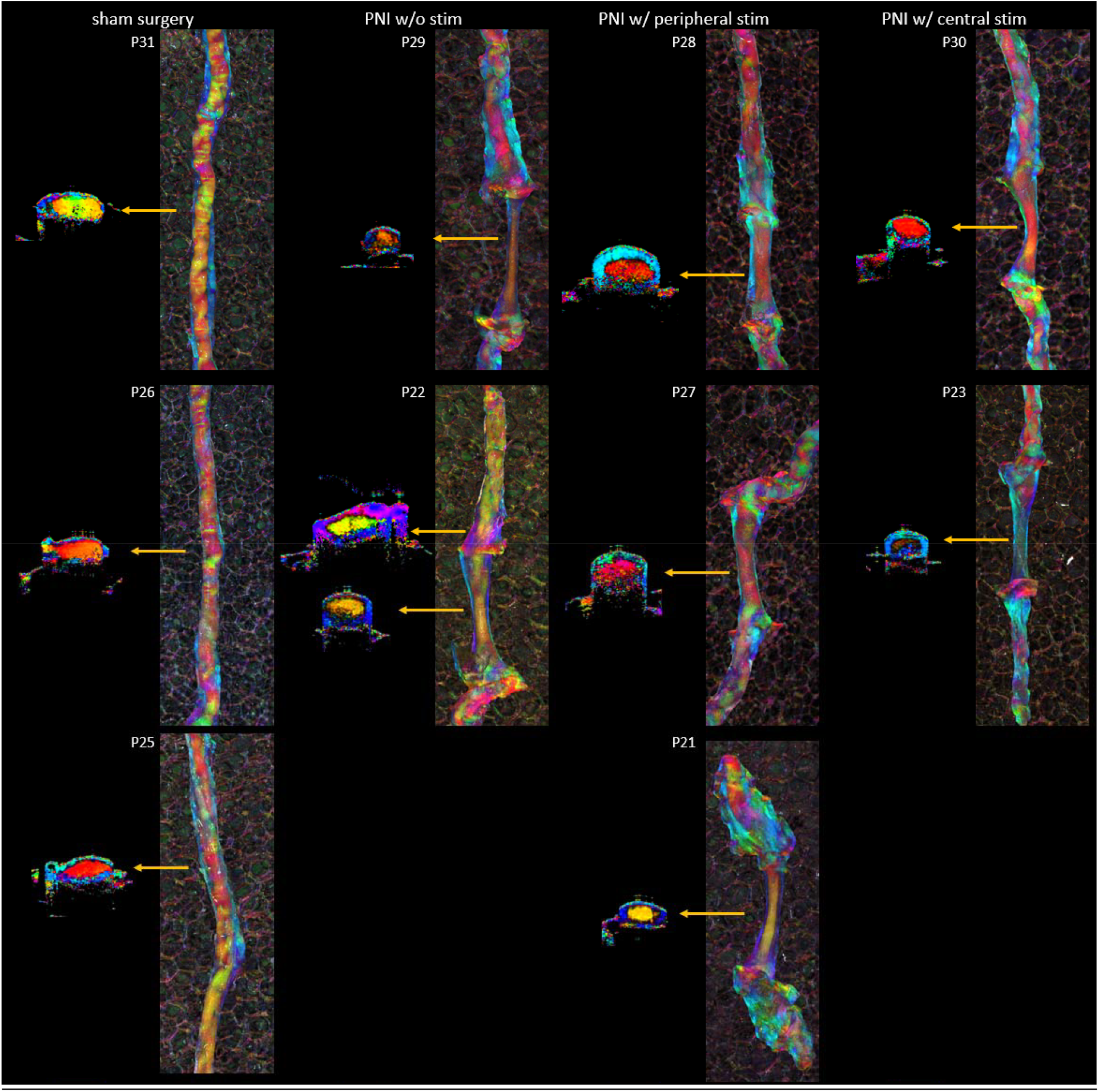
PS-OCT images from 10 representative samples from the Sham, No Stim, Peripheral Stim and Trigeminal Stim groups.

### PS-OCT Imaging of Regenerating Nerves

_____PS-OCT was used to assess the quality of nerve regeneration following rehabilitation and therapeutic approaches. In contrast to traditional histological measures, PS-OCT provides a non-destructive technique to assess nerve structure, including the degree of myelination, both at the site of injury/regeneration and at locations further away. Examination of nerve samples from the historical cohort demonstrated wide variation between the size and structure of nerve samples near the site of injury and subsequent regeneration. Qualitatively, injured nerves had smaller cross sections compared to the sham group, with the poorest regeneration being seen in the group that received rehabilitation alone. In contrast, samples from rats that received both rehabilitation and peripheral stimulation showed larger cross sections with remarkable center/surround organization indicative of regenerating axons. Here, a consistent color in the center of the nerve that contrasts with the color of the epineurium suggests longitudinally aligned myelinated fibers.

## Discussion

Electrical stimulation continues to increase in popularity as a possible avenue to augment the functional outcomes of injured peripheral nerves. Intraoperative electrical stimulation at the site of peripheral nerve injury has been demonstrated to synchronize axonal regeneration across the nerve gap, improving functional outcomes.^9^ This mechanism is thought to enable a synchronous stimulus for the injured axons and supporting cells, enabling a switch to the regenerative mode at the same time point. By a separate and much less studied mechanism, cranial nerve stimulation during the rehabilitation period has shown to augment neuroplasticity of the cerebral cortex, improving functional outcomes.^17^ Throughout rehabilitation, it is hypothesized that this therapy enables a remapping of neurons to accelerate a relearned task thereby restoring function at an earlier time point.

Comparing the functional motor data of the various electrical stimulation groups illustrates a return to baseline function of the combined PS+TNS cohort faster than the PS only group by 1 week and TNS only group by 3 weeks (Figure 3). This improvement supports the hypothesis that the PS and TNS therapies both improve the rate of functional recovery of peripheral nerves through different complimentary mechanisms. PS improves peripheral nerve regeneration, allowing a synchronized regenerative process, while the TNS impacts the subject’s nervous system plasticity at a central level. These modalities may be stacked to reap the benefit of both simultaneously.

Although the functional motor data provides evidence of accelerated return to baseline, the sensory data cannot draw the same conclusions. This is due to the lack of demonstrated sensory injury at the first timepoint of rehabilitation (Figure 4). It would be expected for the PS+TNS cohort to mirror the injury pattern and sensory loss of the PS group at this timepoint, as there has not yet been administration of any TNS therapy at the beginning of rehabilitation. Given that there is no significant difference in the first timepoint captured of the PS+TNS cohort, further investigation is necessary to elucidate the effects of sensory return in this cohort.

This experiment demonstrated the improved functional motor outcomes of the combined PS+TNS cohort of rodents in a peripheral nerve injury model when compared to either of the electrical stimulation therapies alone. The previous investigation completed by Nicksic et al. demonstrated the superior recovery of the injured peripheral nerve when using electrical stimulation (either PS or TNS) compared to no electrical stimulation. This study further demonstrated the additive benefit of combining these two therapies to reap the benefit of the therapies simultaneously. With the growing body of evidence showing an improvement of outcomes when employing electrical stimulation in injured peripheral nerves, we have demonstrated that the additive effect of PS+TNS greatly decreases the rehabilitation time required to return to baseline functional motor status, while functional sensory status remains to be demonstrated. Given it’s non-invasive nature and the lack of side effects reported in clinical trials for FDA approved indications, TNS represents an exciting adjunct therapy for peripheral nerve injury that can easily be administered in conjunction with PNS to improve functional outcomes beyond what is capable with either stimulation modality in isolation.

### Limitations

The completion of the PS+TNS cohort reported herein occurred following the completion of the previous four cohorts may introduce bias in the rodents used and instrumentation handling by a different lead investigators (SCE and PJN). This was most clearly demonstrated in the von Frey sensory assessments, which yielded significantly different results across the experiments. In addition, lack of randomization of this cohort’s rodents with the previous four cohorts following initial training may lead to bias of the PS+TNS cohort when compared to the previous cohorts. This lack of randomization was minimized as much as possible with the use of the same protocol and same species and strain of animals, but differences in initial baseline sensory assessments may be due to these confounding variables. This study utilizes an invasive method for delivering trigeminal nerve stimulation that is then used to infer non-invasive applicability in humans. Invasive methodologies such as this are required in rodent models as a means to ensuring consistent delivery of therapeutic stimulus over an extended period of time.

## Conclusion

Combined electrical stimulation therapy with intraoperative peripheral nerve stimulation and trigeminal nerve stimulation demonstrated a faster return to baseline functional status in motor testing in our rodent peripheral nerve injury model. Moreover, functional sensory recovery returned to baseline pre-injury levels at the first measurement timepoint, indicating an accelerated recovery compared to control. Intraoperative peripheral nerve stimulation and trigeminal nerve stimulation have shown to improve the functional recovery of injured peripheral nerves through separately acting, yet additive, mechanisms. When used simultaneously, these modalities have shown to build upon each other to deliver faster motor functional regeneration in injured peripheral nerves.

## Acknowledgements

The authors would like to thank the University of Wisconsin School of Medicine and Public Health ICTR-Shapiro Medical Student Research Fellowship for supporting SCE during his research year.

## Author Contributions

SCE contributed to study design, data collection, statistical analysis, manuscript drafting and manuscript revision.

PJN contributed to study design and manuscript revision.

DTD contributed to study design, data collection, and manuscript revision.

AJ contributed to data collection.

ECS contributed to data collection and manuscript revision.

WZ contributed to study design, data collection, and manuscript revision.

AJS contributed to study design, statistical analysis, and manuscript revision.

SOP contributed to study design, statistical analysis, and manuscript revision.

AMD contributed to study design, statistical analysis, and manuscript revision.

## Data Availability Statement

All data to support these findings are available from the corresponding author upon request.

